# Energetics and behavior during predation in wild, schooling white mullet (*Mugil curema*)

**DOI:** 10.64898/2026.04.02.716113

**Authors:** Ishani Mukherjee, James C. Liao

## Abstract

Although predation is a major driver of group living across taxa and the antipredator benefits of grouping are well established, the energetic costs experienced by groups under predation remain largely unexplored. In the current study, we use wild, white mullet (*Mugil curema,* Valenciennes 1836), to provide the first real-time quantification of the energetic cost of escape in schooling fish using intermittent, closed-loop respirometry. We found that small groups exposed to predators showed a 53.8% increase in their organismal metabolic rate (MO_2_) as compared to groups without predator exposure. When we evaluated antipredator behaviors such as escape response, group cohesion, and displacement of the group centroid, we found a positive correlation to energetic costs. We then investigated whether escape responses are socially modulated by comparing the energetic costs of escape across solitary individuals, solitary individuals with visual access to a group, and groups. We found that escape frequency and energetic costs to predation were comparable across social contexts, indicating that escape may be an intrinsic survival response independent of cues from group members. Furthermore, we found that fish exposed to predators showed markedly reduced feeding, suggesting that predation constrains energy acquisition in addition to imposing direct energetic costs. Our results provide the first direct quantification of the energetic costs of escape in a schooling fish, offering new insights into the physiological trade-offs underlying collective antipredator defenses.

## Introduction

Predation is among the most powerful selective pressures shaping the behavior, physiology, and life-history traits of animals across ecosystems (Lima and Dill, 1990; Dawkins and Krebs, 1997; Caro, 2005; Sih et al., 2010; Herbert-Read et al., 2017; Sabal et al., 2021; Jermacz et al., 2022). Predator pressure not only drives anti-predator strategies but also shape changes in mating strategies, foraging tactics, and social organization in prey species (Sorato et al., 2012; Wilgers et al., 2014; Herbert-Read et al., 2017). One of the most widespread adaptive responses to predation is group living, which offers multiple antipredator benefits. Individuals in groups experience reduced per capita predation risk through risk dilution, collective vigilance, and predator confusion (Foster and Treherne, 1981; Beauchamp et al., 2012; Beauchamp, 2013; Lehtonen and Jaatinen, 2016). Prey living in groups often exhibit striking collective behaviors when confronted with predators, including synchronized maneuvers such as rapid turns or dives (Beauchamp, 2012; Herbert-Read et al., 2015; Doran et al., 2022; Bierbach et al., 2025), and in some species, alarm signaling or mobbing behaviors that warn conspecifics or actively deter predators (Zimmermann and Curio, 1988; Radford and Ridley, 2006). At the individual level, a key component of predator avoidance across taxa is the execution of escape responses, which can determine survival during encounters (Domenici and Blake, 1997; Blumstein et al., 2016). A striking example of predator evasion are saltwater schooling fish that undertake migrations spanning hundreds of miles while exposed to continuous predation from jacks, tarpons and dolphins (Furey et al., 2018; Luo et al., 2020).

Time-averaged energetic costs such as daily metabolic expenditure associated with activities like migration, foraging and reproduction have been used to estimate overall energy budgets across species (Behrens et al., 2006; Johansen et al., 2020; Soriano-Redondo et al., 2023). While these integrative measures provide valuable insights into how animals allocate energy over extended periods, they obscure the brief, high-intensity bursts of metabolic effort that occur during rapid events such as predator evasion, leaving the immediate energetic costs of these responses largely unresolved. Quantifying these costs in real time is critical because the energetic investment in predator evasion directly constrains the aerobic capacity available for other essential activities such as migration, growth, and reproduction (Rosenfeld et al., 2015; McBride et al., 2015). In this study, we provide the first quantification of the energetic costs incurred *during* escape in a schooling fish, offering new insight into the physiological trade-offs underlying collective antipredator defenses.

Escape events demand rapid mobilization of metabolic capacity. Because escape responses are brief but intense, they require the rapid mobilization of metabolic resources and draw directly on an individual’s available physiological capacity. This capacity is commonly described in terms of aerobic scope, the difference between maximum and resting oxygen consumption. Aerobic scope defines the metabolic capacity available for functions beyond maintenance and is a key determinant of fitness (Clark et al., 2013). Elevated predation risk, through increased escape-related energy expenditure, can reduce aerobic scope and limit energy allocation to growth, reproduction, and foraging. Understanding how predation pressure interacts with aerobic capacity is therefore essential for interpreting both the immediate and longer-term energetic consequences of predator exposure. Under high or continuous predation, fish schools may also rely on their anaerobic energy reserves to escape, which is later replenished through aerobic respiration. This repayment appears as increased oxygen consumption after predation and is known as excess post-exercise oxygen consumption (EPOC) (Scarabello et al., 1993; Hancock and Gleeson, 2008).

Fish rapidly escape a threat by forming a characteristic C-bend of the body, which is actuated by Mauthner cells in the hindbrain (Eaton et al., 1977; Fetcho, 1991; Domenici and Blake, 1997). This bend is rapidly followed by body straightening that quickly accelerates the fish away from the threat. Other behavioral responses to predators in schooling fish include increasing group size, reducing inter-individual distances and seeking refuge. Fish may also modify intrinsic responses to predators by gaining information on predators cues from group members (Kavaliers and Choleris, 2001; Ferrari and Chivers, 2009; Ferrari et al., 2010, Pinheiro et al., 2024; Mukherjee and Bhat, 2025). For instance, solitary fish can initiate escapes at various angles, in contrast to schooling fish that typically escape in straight and uniform trajectories owing to the spatial constraints imposed by the school (Evans et al., 2019). Compared to solitary fish, fish in groups are more prone to initiate escape when nearby individuals react, leading to highly synchronized maneuvers that increase predator confusion (Domenici and Batty, 1997; Mathiron et al., 2015; Hein et al., 2018). Conversely, solitary fish experience isolation-induced stress and this may impact their response to a predator, sometimes decreasing escape probability or extending reaction latency (Galhardo and Oliveira, 2014). Building on these studies, we ask whether social context reshapes the energetic costs associated with escaping predators. In addition to quantifying the direct metabolic expenditures of escape, we also assess indirect costs of predation by measuring changes in foraging behavior. Together, these approaches provide a comprehensive view of how predation shapes the energy budgets of schooling fish.

White mullet are a schooling marine fish that belong to the family Mugilidae, and play a key trophic role in Florida’s coastal ecosystem as an important food source for many predators (Middlemiss et al., 2018). These include dolphins (Atlantic Bottlenose, *Tursiops truncatus*), predatory birds (osprey, *Pandion haliaetus*) and a large diversity of teleost fishes (i.e. crevalle jack *Caranx hippos*, mangrove snapper *Lutjanus griseus*, bluefish *Pomatomus saltatrix*, red drum *Sciaenops ocellatus*) (Whitfield et al., 2012). White mullets are therefore under strong selective pressure in the wild to form protective schooling formations, culminating in migrations numbering in the hundreds of thousands in the summer (Thomson 1955). In this way, mullet schools are rarely subjected to single attacks from solitary predators. Instead, predation in natural habitats often continuous and occurs in persistent bouts from groups of predators, such as a pod of dolphins (Ridgway et al., 2022). For these reasons, mullet are an ideal model for investigating the energetic costs and social context of predator evasion.

Our goals for this study were to, 1) quantify anti-predator responses and real-time energetic costs of mullet groups using intermittent closed respirometry, 2) determine whether social context modulates escape behavior and energetic costs and 3) assess indirect energetic costs of predation by measuring feeding activity in the presence of live predators. We hypothesize that small groups of mullet responding to predator stimuli will display antipredator behaviors along with increased energetic costs. Furthermore, if escape response in schooling fish is socially driven, we expect individuals in isolation to show higher energetic costs as compared to those within groups. Lastly, we hypothesize that feeding activity will decline in the presence of live predators.

## Methods

### Fish collection and maintenance

Wild schools of white mullet were collected with a cast net in January 2025 from the Matanzas River Inlet in Saint Augustine, Florida, USA. Fish (11.51±0.17 cm standard length, mean ±standard error, 25.13±0.70g body weight mean ±standard error) were housed in four 588-liter rectangular tanks (63.5cm × 122cm × 76cm) continuously supplied with fresh, UV-sterilized filtered seawater via an inflow-outflow system. Water temperature was 20 ± 1°C, and the fish holding room was kept under a 12 h:12 h light: dark cycle. Fish were fed once each day *ad libitum* with commercial food pellets (Autohime C2, Reed Mariculture, California, USA). For our experiments, we tested a total of 155 fish.

### Experiments on behavioral responses and associated energetic costs to predator stimuli

#### (1) Experimental treatments

We quantified escape responses and the associated energetic costs in mullets across different social contexts. We conducted experiments involving: (i) groups of four fish (Group), (ii) solitary individuals (Single), and (iii) solitary individuals tested within a group context (Single + G). The Single + G condition is a control to test potential stress arising from social isolation (in Single treatments) by placing a single fish in a transparent inner tank submerged within a larger tank housing three visibly accessible conspecifics (Figures 1A and 1B). We tested ten groups of four fish. Of these, we exposed five to predator stimuli and five without predator stimuli. Similarly, ten fish were tested individually, with five exposed to predator stimuli and five without. We also tested five fish that were individually exposed to predator stimuli while being in visual contact with a group. In total, we tested 55 fish for respirometry experiments. Control and Predator treatments we carried out in a randomized order.

**Figure 1.**
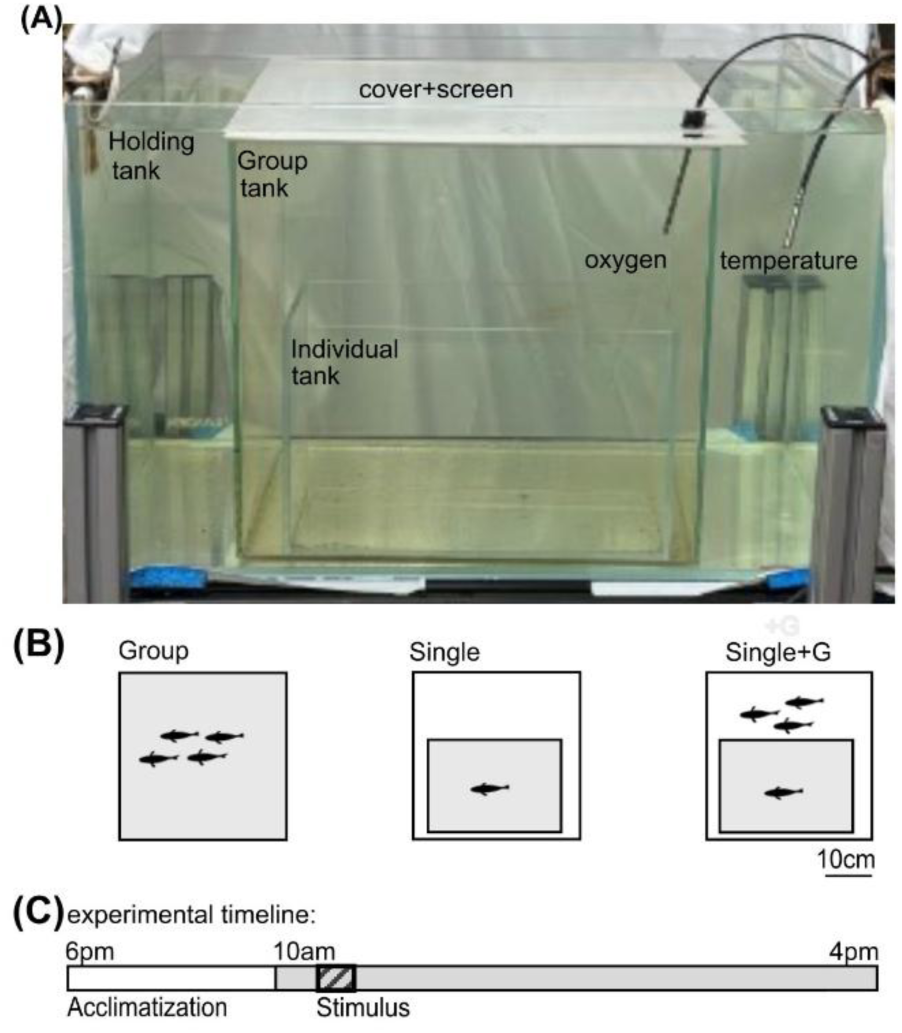
**Setup for recording oxygen consumption and behavior in individual and groups of mullet**. (A) A tank-within-a-tank setup was adopted that consisted of a outermost tank (“Holding” tank; 41 × 41 × 61 cm, 102.5 L), which either contained both a “Group” tank (36 × 36 × 36 cm, 46.65 L) and a smaller “Individual” tank (29 × 20.5 × 15 cm, 8.9 L) nested inside it, or only the Group tank. Filtered sea water fully submerged the inner tank(s). The Group tank was covered with a cover and screen, which enabled the projection of predator stimuli from overhead. If present, the Individual tank was covered with a transparent acrylic lid. A fiber-optic oxygen sensor was inserted through a custom hole in the lid of the test tank and a temperature probe was placed directly in the Holding tank. The entire setup was covered on three sides with white cloth. (B) Three social contexts tested. In the Group condition, four fish were placed together in the Group tank. In the Single condition, a lone test fish was placed in a smaller Individual tank nested within the Group tank. In the Single+G condition, a test fish in the Individual tank could see three conspecifics swimming in the surrounding Group tank. The shaded regions indicate the tank containing experimental fish. The scalebar represents 10 cm. (C) Timeline for experimental trials. Protocol involved overnight acclimatization (6:00 PM to 10:00 AM, white bar), followed by experiments (10:00 AM to 4:00 PM, shaded bar). During experiments, dissolved oxygen concentration and temperature were recorded continuously in 10-minute sessions. Predator treatments occurred in the third 10-minute session (bar with diagonal lines), during which fish were continuously exposed to aversive visual and acoustic stimuli. In control treatments, no stimulus was presented. The schematic is not to scale.

#### (2) Experimental Setup

Our setup consisted of a tank-within-a-tank system to ensure that our measurement tank was free from mixing with atmospheric air, a requisite for respirometry (Lucas et al., 1993). To do this, we placed a 46.65-liter border-less cube tank (herein called the Group Tank, 36cm × 36cm × 36cm) inside a 102.5-liter holding tank (41cm ×41cm ×61cm). The system was filled with filtered seawater, ensuring that the inner tank was fully submerged. We removed air bubbles (if any) from the inner tank. To prevent water exchange between tanks, we then sealed an acrylic cover over the inner tank with a thin layer of dental wax. To carry out Single and Single+G trials, we nested a third tank (Individual Tank, 8.9-liter, 29cm×20.5cm×15 cm). Lower tank volume in the latter treatments ensured a consistent fish mass-to-water volume ratio of ∼1:1000 across trials. Our pilot experiments revealed that these volumes ensured sufficient sensitivity to detect rapid oxygen depletion by this species and body size. In Single+G trials, we placed one fish in the innermost tank and three others in the outer group tank. All tanks in the nested tank system were borderless and transparent, allowing visual access between the inner and outer tanks. We covered the tank-within-a-tank system on three sides with white cloth to block external visual cues, leaving one side open for video recording.

We mounted the tank system on a custom 80/20 aluminum frame (80/20 Inc., Columbia City, IN, USA). To project a looming stimulus, an overhead projector (Epson BrightLink 696Ui; 1920 × 1200, 60 Hz; Epson America, Los Alamitos, CA, USA) was placed above the acrylic tank lid, coated with projection paint (Goo Projector Screen Paint, Alternative Screen Solutions, MI, USA). We placed a portable waterproof speaker (EBODA Speakers, Seattle, USA) adjacent to the tank system to play dolphin vocalizations. We threaded a fiber-optic optical oxygen sensor probe (Wiltrox system, Loligo Systems, Viborg, Denmark) through a fitted hole in the acrylic tank cover. We placed a temperature probe in the outer tank. We used AutoResp v3 (Loligo Systems, Viborg, Denmark) to measure oxygen saturation and temperature every second. We recorded fish movements using two Basler Pylon cameras (Basler AG, Ahrensburg, Germany; 100 frames per second, 904 x 904 pixel resolution), one lateral and one directed at a 45° mirror below the tank for ventral view. We used Streampix (NorPix, Quebec, Canada) to synchronize recordings. All experiments were performed with tanks at room temperature (20°C±1°C).

#### (3) Experimental protocol

Fish were transferred into the inner, experimental tank for overnight acclimation before the start of each experiment. For trials involving individuals tested within a group context, three individuals were present in the outer tank. During this period, the tank was aerated using an air stone bubbler (Marina Long Airstone, Rolf C. Hagen (USA) Corp., Massachusetts, United States). Experiments started at 10:00 am EST the following day, when we removed the bubbler and carefully sealed the lid using a thin, leak-resistant gasket of dental wax. 10 minutes after sealing, we started the trials. Oxygen level and temperature was monitored inside the sealed tank every 1s (Loligo Systems, Viborg, Denmark). If oxygen levels fell below 80%, we paused the trial, opened the tank and aerated until saturation returned to 100%. Organismal metabolic rate (MO_2_) was recorded every 10 minutes for the duration of the experiments, which lasted 5.5 hours.

In half of the trials, we introduced continuous predatory stimuli for 10 minutes, beginning 20 minutes after the start of MO₂ measurements (during the third MO₂ measurement session) (Figure 1C). The third 10-minute session was selected to apply the predatory stimulus to disentangle elevated initial MO₂ values (typically observed due to higher baseline activity earlier in the day) from the energetic demands induced by predation. To mimic the sustained natural predation faced by mullet schools, our predatory stimuli consisted of simultaneously presenting predatory dolphin vocalizations (Ridgway et al., 2022) and a visual looming stimulus. We alternately used two loom sizes: one expanded to cover 2.71% of the screen area and the other expanded to cover 100% of the screen. Both looms expanded linearly for 1s and were played for 10s and were alternated throughout the entire 10-minute predator session to minimize habituation and maintain strong escape. We recorded both ventral and lateral video recordings during the predator stimuli.

We next measured the maximum rate of oxygen consumption for individuals (e.g. maximum aerobic metabolic rate, MMR). After the main trials, we let the fish rest for 30 minutes. We then subjected the fish to a standardized chase protocol (Killen et al., 2017), which involved chasing test fish with a hand net and gently touching their caudal fin for 3 minutes to induce exhaustive activity. Immediately after, we sealed the tank and recorded the oxygen consumption. The resulting value represented the upper physiological limit of aerobic metabolism. Following each trial, we returned fish to a separate housing tank to ensure that all individuals were subjected to the experimental procedure only once.

### Foraging behavior in the presence of live predators

We quantified foraging behavior in the presence live predators by placing ten mullet in a 588-liter tank (63.5cm × 122cm × 76cm) containing a native fish predator (mangrove snapper, *Lutjanus griseus,* L= 20.28 ±0.57cm; mean standard length ± standard error, Figure 5A) contained in a cage (42cm × 36cm × 21cm). These large tank experiments (water volume 12.6 times that of the respirometry tank) extend the scope of the study to more natural conditions. We placed food pellets onto a feeding dish at the tank bottom and recorded foraging behavior from overhead using a GoPro Hero 8 camera (120 frames per second, 1080 x 1920 pixel resolution, GoPro Inc, California, USA). We tested ten groups in total. Of these, we exposed five groups to live predators (Live Predator treatment) and five to an empty predator cage (Control). We tested 100 fish for these foraging behavior experiments. We carried out Control and Live Predator treatments in a randomized order. After testing, fish were transferred to a separate tank to ensure no reuse. We manually counted instances of foraging, defined as when an individual swam onto the food dish and paused with its head tilted down, in a process that was blind to the presence of predators.

## Data Preparation

To assess group-level behavioral responses to predator stimuli, we quantified a range of anti-predator behaviors. We used lateral-view videos to manually count escape events and note the duration of each escape event. An escape was an all or none behavior involving a rapid fast-start swimming maneuver that allowed the fish to quickly accelerate, change direction, and increase distance from the threat (Domenici and Hale, 2019). To estimate whether individuals moved away from the loom towards the tank bottom or corners, we analyzed six randomized videoframes from lateral-view video for each group in Fiji (Schindelin et al., 2012), and quantified the proportion of the group located beyond 3.5 BL from the loom, a threshold based on the loom’s position relative to the tank bottom corners.

We tracked individuals in groups from ventral view recordings using DeepLabCut (Mathis et al., 2018), and estimated group characteristics based on the output trajectories. We calculated the total displacement of the group centroid for the first 45000 frames to assess the movement of entire group in the presence and absence of predator stimuli, for each condition. This involved first determining the group centroid (center point of polygon made by connecting position of all individuals) from individual trajectories in each frame, followed by calculating its total displacement in body lengths (BL). To visualize group cohesion, we generated a heatmap plotting the relative positions of all neighboring fish with respect to each focal fish, normalized to BL. Contour lines enclosed the regions occupied by 25%, 50%, and 75% of the group. To statistically compare group cohesion between treatments, we calculated mean nearest-neighbor distance (NND) or the mean distance between all 4 individuals across all frames.

We calculated energetic costs from measurements of the organismal metabolic rate (MO₂), based on the slope of oxygen decline over a 10-minute interval. To estimate the overall behavioral responses of a group, we ranked several key metrics: (i) escape duration, (ii) displacement of centroid and (iii) mean nearest-neighbor distance (NND) low to high. A ranking summary generated a Response Score, where higher scores indicated stronger behavioral responses. We then quantified the correlation (Spearman’s rank correlation) between MO₂ and Response Score.

To assess the costs experienced by mullets in the presence of live predators, we quantified foraging in presence (and absence) of a predator by manually counting the number of foraging events over a 25-minute period from video recordings.

## Statistical Analysis

We performed all statistical analyses using R Studio (v2024.12.0) (R Core Team, 2024). We used Wilcoxon rank-sum tests (unpaired) to compare escape count, escape duration, nearest-neighbor distance (NND), displacement of group centroid, proportion of individuals beyond 3.5 BL from the loom, MO₂ and the number of bites at food between Predator and Control conditions in groups and/or individuals. For performing multiple comparisons (escape behavior and MO_2_ across Groups, Single and Single+G) we used Wilcoxon rank-sum tests with Bonferroni correction . We used Spearman’s rank correlation to correlate MO₂ to response scores. Given the sample size (n = 5 groups per condition), we employed the Wilcoxon rank-sum test (unpaired) and Spearman’s correlation, both of which are non-parametric methods that do not require assumptions of normality. All tests were two-tailed, and a *p*-value threshold of <0.05 was considered statistically significant. All reported values are presented as means ± standard errors.

## Results

### Behavioral responses and associated energetic costs to predator stimuli

In the presence of predator stimuli, individuals within a group exhibited escape responses (Figure 2A). Both the number of escape events and their durations were significantly higher in groups exposed to predator stimuli compared to controls: groups exposed to predator stimuli exhibited 12.60 ± 0.38 escapes versus 1.40 ± 1.20 escapes in controls (Unpaired Wilcoxon Test: W = 23, p= 0.03, Figure 2B), and escape durations of 16.8 ± 5.58s compared to 1.6± 1.43 s in controls (Unpaired Wilcoxon Test: W = 23, p= 0.03, Figure 2C). Also, individuals within groups exhibited avoidance behavior by aggregating near the tank corners. A significantly greater proportion maintained a distance of at least 3.5 body lengths (BL) from the top center of the tank during loom presentations (0.34 ± 0.08), compared to no-loom trials (0.07 ± 0.04) (Wilcoxon test results: W = 23.5, p = 0.02; Figure 2D).

**Figure 2.**
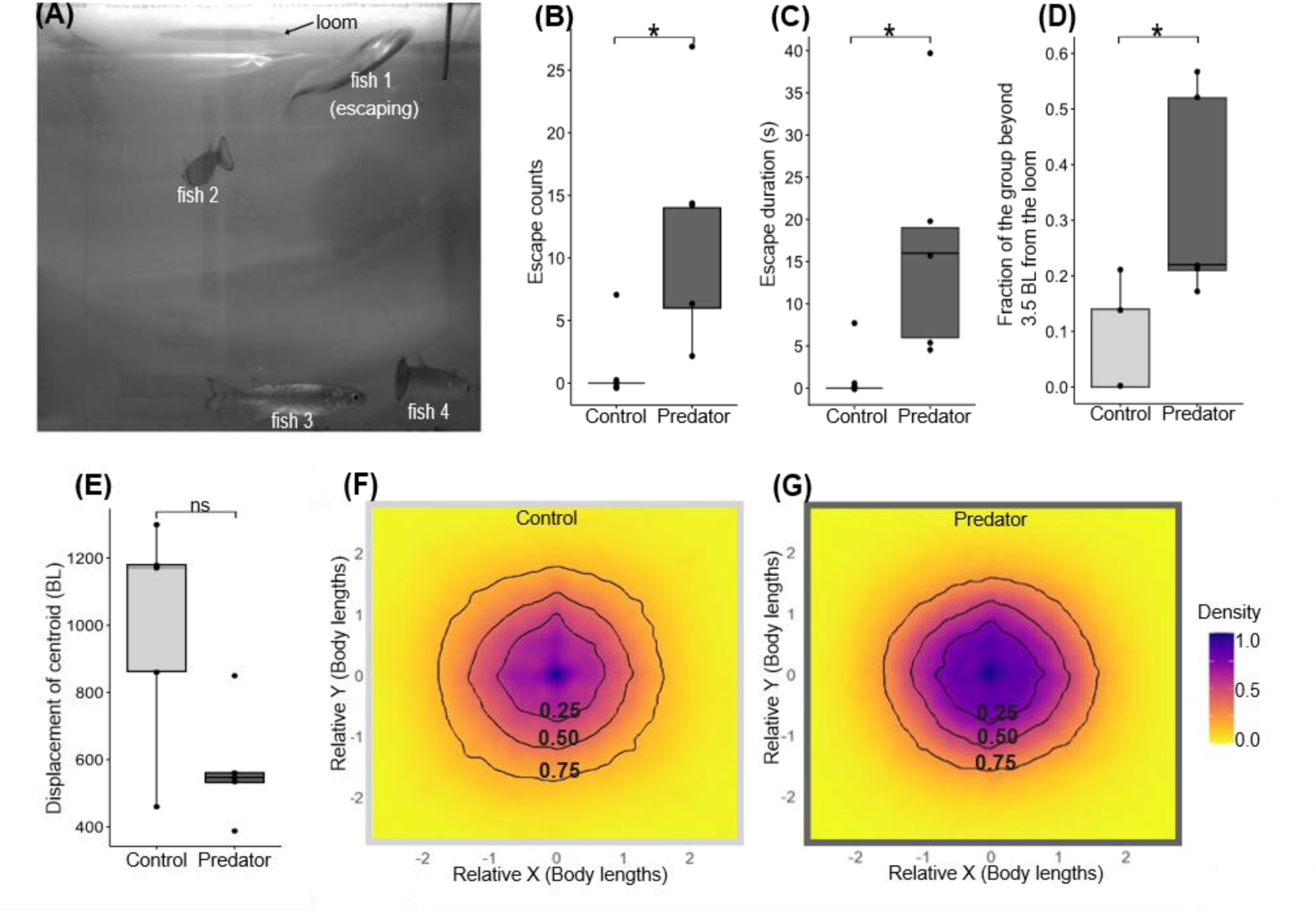
Predator stimuli elicit changes in group behavior. (A) Representative still image from video recording showing fish responses to the looming stimulus. Fish 1 is actively escaping, while Fish 2-4 are not. The expanding looming stimulus is visible at the top of the frame. (B-E) Box and whisker plots showing behavior under Control (light gray) and Predator (dark gray) treatments. (B) Escape counts, (C) Escape duration (seconds) and (D) Fraction of the group beyond 3.5 body lengths (BL) from the loom stimulus were significantly greater in Predator treatments as compared to Control treatments. (E) Displacement of the group centroid (BL) was comparable between Predator and Control treatments. Data points for each school are represented as dots. n.s. indicates no significant difference and * indicate statistically significant difference (p<0.05). (F, G) Heatmaps depicting positions of individuals relative to a focal fish at the center (0,0) for (F) Control and (G) Predator treatments. Black contour lines indicate portion of group proportion (25%, 50%, 75%) around focal fish. The color bar represents the density of fish.

The total displacement of the group centroid was not significantly different between predator and control groups (Predator: 575.11±66.90 BL, Control: 995.61±135.69 BL, Wilcoxon test results: W=4. P=0.09, Figure 2E). Groups were more cohesive under predation risk (Figure 2E & Figure 2F), a pattern further supported by nearest-neighbor distance (NND) analyses. NND values were significantly lower during predator exposure (0.97 ± 0.03 BL) compared to the no-exposure condition (1.12 ± 0.04 BL; Unpaired Wilcoxon test: W = 3, p = 0.05; Figure S1) indicating increased group compactness as a potential anti-predator strategy.

Mean MO₂ values (± SE) were similar between predator and control groups before and after predator cue exposure (Figure 3A). When predator stimuli were presented, MO₂ was significantly higher in the predator group (585.69 ± 38.95 mg O_2_ kg^-1^hr ^-1^) compared to the control group (380.79 ± 46.69 mg O_2_ kg^-1^hr ^-1^; Unpaired Wilcoxon test: W = 23, p = 0.03; Figure 3B). The maximum metabolic rate (MMR) was 766.58 ± 68.04 mg O_2_ kg^-1^hr ^-1^. Notably, two schools exposed to predator stimuli exhibited elevated MO₂ values (of 668.88 and 677.2mg O_2_ kg^-1^hr ^-1^) approaching the species’ MMR. Overall, the energy use (mean MO_2_ of groups) under predator exposure accounted for approximately 76.4% of the total metabolic capacity. The additional energy demand due to predator presence reflects a substantial metabolic cost, leaving 23.6% of the aerobic scope unused under threat conditions (Figure 3C).

**Figure 3.**
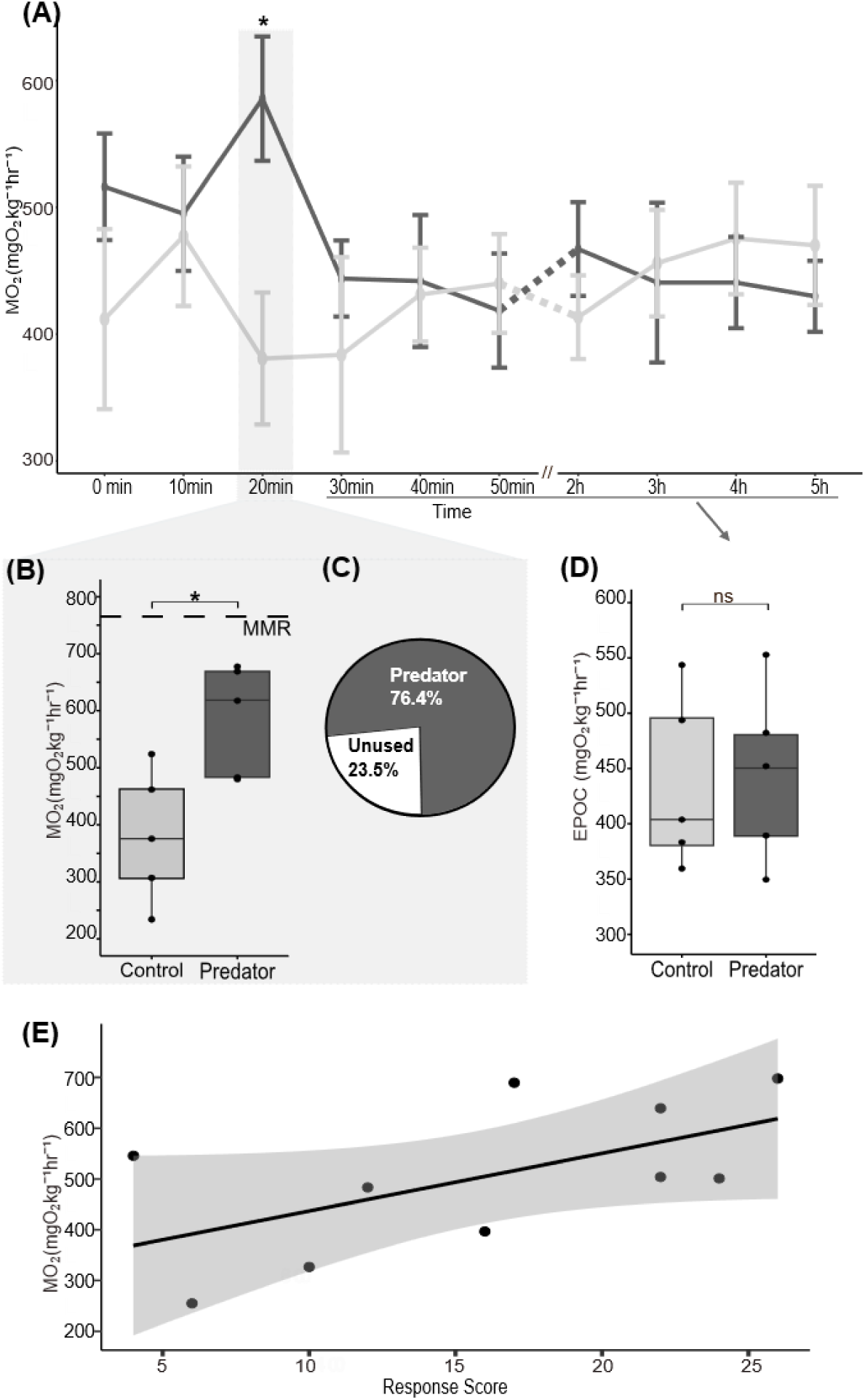
Energetic costs of predation in groups. (A) Temporal profile of organismal metabolic rate or MO₂ (mg O₂ kg⁻¹ h⁻¹) in Control (light gray) and Predator (dark gray) treatments. Values represent the mean ± standard error of five groups. At the 20 minute mark in the third session (shaded), groups in the Predator treatment were exposed to predator stimuli (control groups were not exposed to predator stimuli). (B) Box and whisker plots showing that predator-exposed fish have significantly higher MO₂ (dark gray) than Control fish (light gray, p < 0.05). Dashed line represents the maximum metabolic rate (MMR). (C) Pie chart showing the proportion of MMR utilized during predator exposure (dark gray). (D) Mean MO_2_ after predator stimuli (dark gray) is similar to Control (light gray). Each dot denotes escape count or MO_2_ value for a single school. n.s. indicates no significant difference and * indicates a statistically significant difference. (E) Scatterplot showing a positive correlation between MO₂ and Response Score (see Methods) for all 10 trials performed.

The energetic costs to predator encounters appeared confined to the period of exposure, as no sustained elevation in MO_2_ (Excess Post-exercise Oxygen Consumption or EPOC) was observed in the predator group (Figure 3D). Specifically, the mean MO_2_ from after the predator session to the end of the experiment was 448.26± 32.09 mg O_2_ kg^-1^hr ^-1^ and 454.12± 32.67 mg O_2_ kg^-1^hr ^-1^for predator and control groups respectively (Unpaired Wilcoxon test: W = 10.5, p = 0.75; Figure 3D). A Spearman’s rank correlation revealed a positive but non-significant association between Response Score and MO₂ values (ρ = 0.58, *S* = 69.71, p = 0.08; Figure 3E).

### Escape responses and associated energetic costs across social contexts

Individuals in a Group did not always escape in synchrony. Among the 28 escape events observed across the five predator-exposed groups, all four individuals escaped together in only 9 out of 28 events. In the remaining events (19 out of 28), partial group escapes were observed (Figure S2). We found comparable escape count per individuals among Single (4.80±1.90 escapes), Single+G (3.00±0.48 escapes) and Groups (3.15±0.95 escapes; Unpaired Wilcoxon test with Bonferroni correction results- Single vs. Group: W = 9, p =1; Single+G vs Single: W=11, p=1; Single+G vs Group: W=12, p=1, Figure 4A). Similarly, the MO_2_ during exposure to predator stimuli was comparable across the individuals as Single (630.30±120.77mg O_2_ kg^-1^hr ^-1^), Single+G (653.32±145.42 88mg O_2_ kg^-1^hr ^-1^) and Group (585.68±38.94mg O_2_ kg^-1^hr ^-1^, Wilcoxon test with Bonferroni correction results- Single vs. Group: W = 13, p =1; Single+G vs Single: W=12, p=1; Single+G vs Group: W=12, p=1, Figure 4B). MO₂ levels were elevated even in the absence of predator stimuli for the Single treatment (472.91 ± 32.55 mg O₂ kg⁻¹ hr⁻¹) and did not differ significantly from levels recorded during predator exposure (630.30 ± 120.77 mg O₂ kg⁻¹ hr⁻¹; Wilcoxon rank-sum test: W = 17, p = 0.42, Figure 4C).

**Figure 4.**
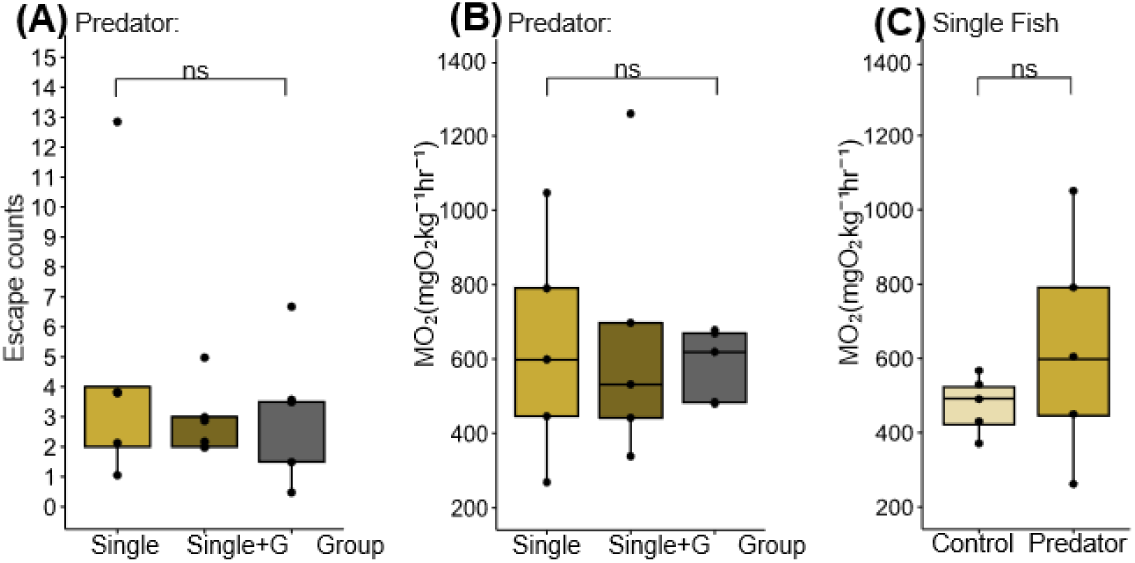
Social context does not impact escape behavior and energetic costs imposed by predator stimuli. Box and whisker plots showing: **(A)** Escape counts and (**B)** MO₂ in predator treatments across social contexts: Single (mustard), Single+G (dark mustard), and Group (dark gray); **(C)** MO2 in Single fish across Control (light mustard) and Predator (mustard) treatments Each dot denotes escape count or MO2 value for a single school. Points outside the whiskers represent outliers. n.s. indicates no significant difference across treatments.

**Figure 5.**
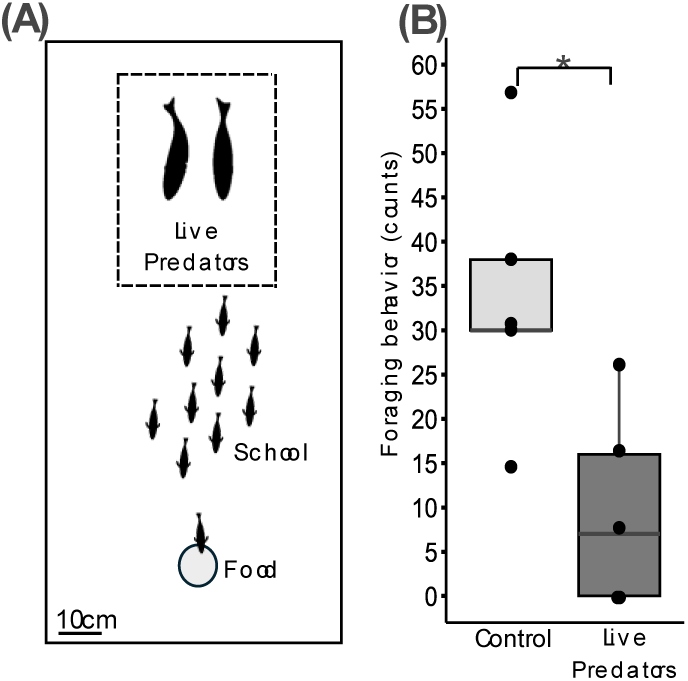
**Indirect energetic costs in the presence of a predator**. (A) Experimental setup showing a predator cage (comprising 2 live predators), a school of test fish (10 mullet), and a food source. The scalebar represents 10cm. (B) Foraging behavior (counts) was significantly lower in live predator-exposed groups than in controls (p < 0.05). Each dot denotes foraging behavior count for a single school. * indicates a statistically significant difference.

### Feeding suppression by live predators

To account for indirect energetic costs of predation risk, we exposed mullet to live predators (e.g. mangrove snapper). Exposure to snappers led to a ∼71% suppression in foraging activity (9.80 ± 4.47 bites) relative to control groups (33.80 ± 6.25 bites; Wilcoxon rank-sum test: W = 2, p = 0.03; Figure 5B).

## Discussion

Our study is the first to measure both the direct, energetic and indirect, behavioral costs of a schooling fish escaping from a predator. Groups of wild mullet quickly distance themselves from predator stimuli by performing rapid escapes with heightened school cohesion. Interestingly, we found that escape energetics and behavior were similar for individual fish regardless of whether they were solitary or with a group. We also observed that groups exposed to live predators suppressed foraging activity. By quantifying the metabolic cost of anti-predator behavior in schooling fish, this study considers physiological, behavioral, and ecological perspectives to better understand the nature of predator-prey interactions.

### Behavioral responses and associated energetic costs to predator stimuli

A group of mullet, as well as individuals within the group, showed pronounced anti-predator behaviors when exposed to predator cues. Individuals within groups escaped to continuous loom stimuli and auditory cues, similar to other fish species (Lefrançois et al., 2005; Marras and Domenici, 2013; Domenici and Hale, 2019; Rodriguez-Pinto et al., 2024), and positioned themselves away from the loom source. Similar spatial avoidance to projected loom stimuli has been found in natural marine habitats, where coral reef fishes adjust their escape trajectories to steer away from threats and swim towards shelter (Hein et al., 2018). We found no significant change in centroid displacement of groups in the presence of a predator (a metric that characterizes collective behavior, Herbert-Read et al., 2013; Jolles et al., 2017), which suggests that centroid-based metrics provide only an estimate of school movement, and fails to capture critical aspects of antipredator behavior such as fine-scale individual escape maneuvers. For instance, if individuals escape in opposite directions, centroid displacement will resemble the lack of response characterizing the pre-predator state. As a result, centroid-based measures may overlook within-group movement variation. This limitation could account for the lack of difference between control and predator-exposed schools. Under predator stimuli, heightened cohesion may not only confuse predators and dilute individual risk but also enhance the speed and precision of collective escape responses (Foster and Treherne, 1981; Lima, 1995; Herbert-Read et al., 2017). Similar to our study, other schooling species such as golden shiners (*Notemigonus crysoleucas*), guppies (*Poecilia reticulata*), fathead minnows (*Pimephales promelas*) and zebrafish (*Danio rerio*) also show greater cohesion upon exposure to predation (Johannes 1993; Chivers et al., 1995; Herbert-Read et al., 2017; Ioannou et al., 2017, Mukherjee and Bhat, 2024).

Our study demonstrates that the antipredator responses deployed by groups under predation risk come at a substantial energetic cost (e.g. a 53.8% increase in oxygen uptake). Hence, our findings shed light on a fundamental trade-off: while grouping is known to reduce chances of predation, groups continue to bear significant energetic costs under threat. The resulting increase in physiological demand may constrain individual energy budgets, with consequences for growth, reproduction, and longer-term fitness. Interestingly, following overnight acclimatization, oxygen intake was slightly higher in the early part of the experiment than later in the day. This may be because fish are naturally more active in the morning (as reported by Helfman 1981; Bosiger and McCormick, 2014), leading to greater oxygen consumption or due to tank-sealing prior to an experiment (followed by a brief second acclimatization) causes mild stress. Hence, we compared our findings to a control performed on a separate group rather than to pre-predator measurements from the same group.

While a predatory stimulus substantially increased oxygen consumption in mullet, it remained below their maximal metabolic rate (MMR), suggesting that mullet can tolerate prolonged predation. MMR and aerobic scope are related to lifestyle across fish species, with sedentary species showing low values and more athletic species displaying relatively high values (Norin and Clark, 2016; Norin and Metcalfe, 2019; Jermacz et al., 2020; Fu et al., 2022). This framework is consistent with our results: migratory mullet possesses high MMR and aerobic scope, reflecting their active lifestyle. Likewise, our results indicate no significant energetic carry-over effects of predation: The MO₂ measured from the end of the predator session to the end of the experiment did not differ between predator and control groups. This result is contrary to what has been found in other marine fishes. For instance, striped surfperch (*Embiotoca lateralis*) exhibit elevated EPOC (excess post-exercise oxygen consumption) following prolonged swimming under flow (Cordero et al., 2019). Atlantic salmon (*Salmo salar*) show similar post-exercise costs after exhaustive chase trials (Zhang et al., 2018). Closely related, golden gray mullet (*Chelon auratus*) exhibits energetic costs after predator attacks, with faster-responding individuals showing higher EPOC, highlighting a trade-off between vigilance and post-exercise metabolic cost (Killen et al., 2015). The elevated EPOC in previous studies likely reflects the use of a mechanical stimulus (e.g. dropping a PVC pipe), designed to mimic a sudden predator strike. In contrast, our aversive stimuli were patterned after the chronic presence of predators, as might be perceived by a fish school under repeated, sustained attack. Thus, the absence of carry over effects, together with the finding that MO_2_ remained below MMR during predation, is consistent with the ecology of wild mullets. Migrating mullet schools face intense predation risk (Richards and Castagna, 1976) and may have evolved to withstand such challenges. Their ability to maintain predation costs below MMR and within aerobic scope, without carry-over effects, may be particularly advantageous in coastal habitats where predator encounters are frequent and unpredictable (Mosman et al., 2023).

Although the relationship between response score and MO₂ was positive, it was not statistically significant. This is likely because antipredator responses are multidimensional, encompassing behaviors such as rapid escapes, cohesion, and subtle postural adjustments, many of which may not be fully captured in our study and within the response score. Furthermore, individuals may differ in the extent to which they elicit antipredator responses and consume oxygen. These considerations suggest that while stronger antipredator responses tend to be associated with higher energetic costs, fully resolving this relationship may require finer-scale behavioral classification and the examination of predator induced changes in physiology in greater details. Similar to our finding, several other fish species such as hammerhead sharks (*Sphyrna lewini*) and southern catfish (*Silurus meridionalis*) show a positive correlation between activity and MO_2_ (Gleiss et al., 2010; Zhang et al., 2010)

### Escape responses and associated energetic costs across social contexts

We expected that schooling would reduce energetic costs due to hydrodynamic wake recapture (Liao, 2022) and social buffering of stress (Culbert et al., 2019). However, the frequency and energetic costs of escapes did not differ for individuals with and without visual access to a school, suggesting that escapes are not strongly influenced by social groups. Our study thus suggests that although grouping may reduce the likelihood of successful predator attacks, this benefit is achieved without increasing energetic costs relative to solitary escape. However, grouping does not appear to confer energetic advantages during predator evasion. This discrepancy likely arises because mullets did not exhibit coordinated escape maneuvers. Instead, individuals fled asynchronously and in divergent directions (as shown in Figure S1) thereby likely disrupting the spatial organization necessary to generate hydrodynamic advantages (Weihs, 1975; Fish and Lauder, 2006). Moreover, the lack of group synchronization during escapes may reflect individual rather than collective decision-making under threat. Given that schooling can provide hydrodynamic advantages under flow (Zhang and Lauder, 2024), we speculate that the presence of water flow may promote more coordinated escape responses among individuals, thereby conferring energetic benefits to solitary fish in ways that were not observed for our fish in still water.

Note that solitary individuals exhibited elevated MO₂ even in the absence of predator stimuli, which may reflect a level of stress associated with isolation. Higher energetic costs and stress due to isolation have also been previously reported across various fish species (Galhardo and Oliveira, 2014; Forsatka et al., 2024; Xu et al., 2024).

### Costs faced by schools facing live predators

The costs of being exposed to predators are not restricted to the immediate energetic demands of escape but also include indirect effects in the form of reduced food uptake. Several fish species decrease their feeding activity in the presence of predators (Des Roches et al., 2021; Shapiro et al., 2021; Mukherjee and Bhat 2025; Ling et al., 2019). Reduced food intake can limit the energy available for sustaining high metabolic performance and growth (Auer et al., 2015; Ling et al., 2019), thereby constraining long-term fitness. While our study quantifies the direct energetic costs associated with heightened oxygen demand during escape behaviors, it also reveals indirect costs arising from behavioral trade-offs such as reduced foraging.

We propose several directions for future work. Measuring latency to escape in response to looming stimuli could provide insights into the responsiveness of wild mullet and may serve as an indirect indicator of the perceived strength of chronic predator stimuli. Also, introducing hydrodynamic flow could test whether predator-induced energetic costs are exacerbated under stress. Finally, investigating neurophysiological correlates, for example through lateral line ablation, could help elucidate the sensory contributions to escape energetics. These approaches would deepen our understanding of how social and environmental factors shape the metabolic consequences of anti-predator behavior in schooling fish. Lastly, investigations quantifying escape behaviors in the wild would be invaluable. In nature, mullets escape predators by fleeing over greater distances and also display behaviors such as leaping out of the water (Peterson, 1976; Whitfield et al., 2012). The confined boundaries of laboratory tanks prohibit a comprehensive understanding of the strategies and costs of predator evasion.

By directly linking escape behavior to metabolic expenditure, our study shows that predator encounters impose substantial yet physiologically moderate energetic costs, and that these costs arise independently of social context. The additional suppression of feeding under predation further underscores that predators influence not only the immediate energetic demands of escape, but also the longer-term balance between energy intake and expenditure. More broadly, our work provides a rare empirical window into the energetic underpinnings of collective behavior and highlights how both direct and indirect metabolic pressures may shape the evolution of sociality, foraging strategies, and risk management across animal taxa.

## Acknowledgements

We would like to thank Glen Greenwald, Peter Meyers, John Perkner, Fan Yang and Zihao Huang for help collecting and maintaining fish.

## Funding

Funding: This work was supported by the National Science Foundation IOS 1856237, IOS 1932707, PHY 2102891 and NSF CMMI 2345913 to J.C.L.

All experimental procedures were approved IUACUC (202200000056)

## Author Contributions

J.C.L., I.M. Conceptualization, J.C.L., I.M. Methodology, J.C.L., I.M. Validation, I.M. Formal analysis, J.C.L, I.M. Investigation, J.C.L. Resources, I.M. Data curation, I.M. Writing - original draft, J.C.L., I.M. Writing - review & editing, J. C.L. Supervision, J.C.L. Project administration, J.C.L. Funding acquisition.

## Data Availability

All data pertaining to this study can be provided upon reasonable request.

## References

1. Auer, S. K., Salin, K., Rudolf, A. M., Anderson, G. J., & Metcalfe, N. B. (2015). The optimal combination of standard metabolic rate and aerobic scope for somatic growth depends on food availability. Funct. Ecol., 29(4), 479–486.

2. Beauchamp, G. (2012). Flock size and density influence speed of escape waves in semipalmated sandpipers. Anim. Behav., 83(4), 1125–1129.

3. Beauchamp, G. (2013). Social predation: how group living benefits predators and prey. London, UK: Elsevier.

4. Beauchamp, G., Alexander, P., & Jovani, R. (2012). Consistent waves of collective vigilance in groups using public information about predation risk. Beh. Ecol., 23(2), 368–374.

5. Behrens, J. W., Præbel, K., & Steffensen, J. F. (2006). Swimming energetics of the Barents Sea capelin (*Mallotus villosus*) during the spawning migration period. J. Exp. Mar. Biol. Ecol. 331(2), 208–216.

6. Bierbach, D., Lukas, J., Gómez-Nava, L., Francisco, F. A., Arias-Rodriguez, L., Krause, S., Korbinian, P., Yunus, S., Pawel, R., & Krause, J. (2025). Collective escape waves provide a generic defence against different avian predators. R. Soc. Open Sci., 12(3), 241055.

7. Blumstein, D. T., Samia, D. S., & Cooper Jr, W. E. (2016). Escape behavior: dynamic decisions and a growing consensus. Curr. Opin. Behav. Sci., 12, 24–29.

8. Bosiger, Y. J., & McCormick, M. I. (2014). Temporal links in daily activity patterns between coral reef predators and their prey. PLoS One, 9(10), e111723.

9. Caro, T. M. (2005). Antipredator defenses in birds and mammals. Chicago, IL: University of Chicago Press.

10. Chivers, D. P., Brown, G. E., & Smith, R. J. F. (1995). Familiarity and shoal cohesion in fathead minnows (*Pimephales promelas*): implications for antipredator behaviour. Can. J. Zool., 73(5), 955–960.

11. Cordero, G. A., Methling, C., Tirsgaard, B., Steffensen, J. F., Domenici, P., & Svendsen, J. C. (2019). Excess postexercise oxygen consumption decreases with swimming duration in a labriform fish: Integrating aerobic and anaerobic metabolism across time. J. Exp. Zool. A Ecol. Integr. Physiol. 331(10), 577–586.

12. Culbert, B. M., Gilmour, K. M., & Balshine, S. (2019). Social buffering of stress in a group-living fish. Proc. R. Soc. B., 286(1910), 20191626.

13. Curio, E. (1978). The adaptive significance of avian mobbing: I. Teleonomic hypotheses and predictions. Z. Tierpsychol., 48(2), 175–183.

14. Dawkins, R., & Krebs, J. R. (1979). Arms races between and within species. Proc. R. Soc. B., 205(1161), 489–511.

15. Des Roches, S., Robinson, R. R., Kinnison, M. T., & Palkovacs, E. P. (2022). The legacy of predator threat shapes prey foraging behaviour. Oecologia, 198(1), 79–89.

16. Domenici, P. (2002). The visually mediated escape response in fish: predicting prey responsiveness and the locomotor behaviour of predators and prey. Mar. Freshw. Behav. Physiol., 35(1-2), 87–110.

17. Domenici, P., & Batty, R. S. (1997). Escape behaviour of solitary herring (*Clupea harengus*) and comparisons with schooling individuals. Mar. Biol., 128(1), 29–38.

18. Domenici, P., & Blake, R. W. (1997). The kinematics and performance of fish fast-start swimming. J. Exp. Biol, 200(8), 1165–1178.

19. Domenici, P., & Hale, M. E. (2019). Escape responses of fish: a review of the diversity in motor control, kinematics and behaviour. J. Exp. Biol., 222(18), jeb166009.

20. Doran, C., Bierbach, D., Lukas, J., Klamser, P., Landgraf, T., Klenz, H., … & Krause, J. (2022). Fish waves as emergent collective antipredator behavior. Curr. Biol., 32(3), 708–714.

21. Eaton, R. C., Bombardieri, R. A., & Meyer, D. L. (1977). The Mauthner-initiated startle response in teleost fish. J. Exp. Biol., 66(1), 65–81.

22. Evans, D. A., Stempel, A. V., Vale, R., & Branco, T. (2019). Cognitive control of escape behaviour. Trends Cogn. Sci., 23(4), 334–348.

23. Ferrari, M. C., & Chivers, D. P. (2009). Latent inhibition of predator recognition by embryonic amphibians. Biol. Lett., 5(2), 160–162.

24. Ferrari, M. C., Wisenden, B. D., & Chivers, D. P. (2010). Chemical ecology of predator–prey interactions in aquatic ecosystems: a review and prospectus. Can. J. Zool., 88(7), 698–724.

25. Fetcho, J. R. (1991). Spinal network of the Mauthner cell (Part 2 of 2). Brain Behav. Evol., 37(5), 307–316.

26. Forsatkar, M. N., Safari, O., & Boiti, C. (2017). Effects of social isolation on growth, stress response, and immunity of zebrafish. Acta Ethol., 20(3), 255–261.

27. Foster, W. A., & Treherne, J. E. (1981). Evidence for the dilution effect in the selfish herd from fish predation on a marine insect. Nature, 293(5832).

28. Furey, N. B., Armstrong, J. B., Beauchamp, D. A., & Hinch, S. G. (2018). Migratory coupling between predators and prey. *Nat*. Ecol. Evol., 2(12), 1846–1853.

29. Fu, S. J., Dong, Y. W., & Killen, S. S. (2022). Aerobic scope in fishes with different lifestyles and across habitats: trade-offs among hypoxia tolerance, swimming performance and digestion. Comp. Biochem. Physiol. A Mol. Integr. Physiol., 272, 111277.

30. Galhardo, L., & Oliveira, R. F. (2014). The effects of social isolation on steroid hormone levels are modulated by previous social status and context in a cichlid fish. Horm. Behav., 65(1), 1–5.

31. Hein, A. M., Gil, M. A., Twomey, C. R., Couzin, I. D., & Levin, S. A. (2018). Conserved behavioral circuits govern high-speed decision-making in wild fish shoals. Proceedings of the National Academy of Sciences, 115(48), 12224–12228.

32. Helfman, G. S. (1981). Twilight activities and temporal structure in a freshwater fish community. Can. J. Fish. Aquat. Sci., 38(11), 1405–1420.

33. Herbert-Read, J. E., Buhl, C., Hu, F., Ward, A. J., & Sumpter, D. J. (2015). Initiation and spread of escape waves within animal groups. Royal Society open science, 2(4), 140355.

34. Herbert-Read, J. E., Krause, S., Morrell, L. J., Schaerf, T. M., Krause, J., & Ward, A. J. W. (2013). The role of individuality in collective group movement. Proc. R. Soc. B. 280(1752), 20122564.

35. Herbert-Read, J. E., Rosén, E., Szorkovszky, A., Ioannou, C. C., Rogell, B., Perna, Ramnarine, I.W., Kotrschal, A., Kolm, N., Krause N., & Sumpter, D. J. (2017). How predation shapes the social interaction rules of shoaling fish. Proc. R. Soc. B. 284(1861), 20171126.

36. Ioannou, C. C., Ramnarine, I. W., & Torney, C. J. (2017). High-predation habitats affect the social dynamics of collective exploration in a shoaling fish. Sci.Adv., 3(5), e1602682.

37. Jermacz, Ł., Kletkiewicz, H., Poznańska-Kakareko, M., Klimiuk, M., & Kobak, J. (2022). Chronic predation risk affects prey escape abilities through behavioral and physiological changes. Behav. Ecol., 33(1), 298–306.

38. Jermacz, Ł., Nowakowska, A., Kletkiewicz, H., & Kobak, J. (2020). Experimental evidence for the adaptive response of aquatic invertebrates to chronic predation risk. Oecologia, 192(2), 341–350.

39. Johannes, M. R. (1993). Prey aggregation is correlated with increased predation pressure in lake fish communities. Can. J. Fish. Aquat. Sci., 50(1), 66–73.

40. Johansen, J. L., Akanyeti, O., & Liao, J. C. (2020). Oxygen consumption of drift-feeding rainbow trout: the energetic tradeoff between locomotion and feeding in flow. J. Exp. Biol, 223(12), jeb220962.

41. Jolles, J. W., Boogert, N. J., Sridhar, V. H., Couzin, I. D., & Manica, A. (2017). Consistent individual differences drive collective behavior and group functioning of schooling fish. Curr. Biol., 27(18), 2862–2868.

42. Kavaliers, M., & Choleris, E. (2001). Antipredator responses and defensive behavior: ecological and ethological approaches for the neurosciences. Neurosci. Biobehav. Rev., 25(7-8), 577–586.

43. Killen, S. S., Reid, D., Marras, S., & Domenici, P. (2015). The interplay between aerobic metabolism and antipredator performance: vigilance is related to recovery rate after exercise. Front. Physiol., 6, 111.

44. Killen, S. S., Norin, T., & Halsey, L. G. (2017). Do method and species lifestyle affect measures of maximum metabolic rate in fishes? J. Fish Biol., 90(3), 1037–1046.

45. Lefrançois, C., Shingles, A., & Domenici, P. (2005). The effect of hypoxia on locomotor performance and behaviour during escape in *Liza aurata*. J. Fish Biol, 67(6), 1711–1729. *Journal of Fish Biology*, *75*(7), 1615-1625.

46. Lehtonen, J., & Jaatinen, K. (2016). Safety in numbers: the dilution effect and other drivers of group life in the face of danger. Behav. Ecol. Sociobiol., 70(4), 449–458.

47. Lima, S. L. (1995). Back to the basics of anti-predatory vigilance: the group-size effect. Anim. Behav., 49(1), 11–20.

48. Lima, S. L., & Dill, L. M. (1990). Behavioral decisions made under the risk of predation: a review and prospectus. Can. J. Zool, 68(4), 619–640.

49. Ling, H., Fu, S. J., & Zeng, L. Q. (2019). Predator stress decreases standard metabolic rate and growth in juvenile crucian carp under changing food availability. Comp. Biochem. Physiol. A Mol. Integr. Physiol., 231, 149–157.

50. Lucas, M. C., Johnstone, A. D. F., & Tang, J. (1993). An annular respirometer for measuring aerobic metabolic rates of large, schooling fishes. J. Exp.Biol., 175(1), 325–331.

51. Luo, J., Ault, J. S., Ungar, B. T., Smith, S. G., Larkin, M. F., Davidson, T. N., Bryan, D.R., Farmer, N.A., Holt, S.A., Alford., A.S., Adams, A.J., Humston, R., Marton, A.S., David, M., Kleppinger, R., Requejo, A., & Robertson, J. (2020). Migrations and movements of Atlantic tarpon revealed by two decades of satellite tagging. Fish Fish., 21(2), 290–318.

52. Mathis, A., Mamidanna, P., Cury, K. M., Abe, T., Murthy, V. N., Mathis, M. W., & Bethge, M. (2018). DeepLabCut: markerless pose estimation of user-defined body parts with deep learning. Nat. Neurosci., 21(9), 1281–1289.

53. Marras, S., & Domenici, P. (2013). Schooling fish under attack are not all equal: some lead, others follow. PLoS One, 8(6), e65784.

54. Mathiron, A. G., Crane, A. L., & Ferrari, M. C. (2015). Individual vs. social learning of predator information in fish: does group size affect learning efficacy? Behav. Ecol. Sociobiol., 69(6), 939–949.

55. McBride, R. S., Somarakis, S., Fitzhugh, G. R., Albert, A., Yaragina, N. A., Wuenschel, M. J., Alonso-Fernández, A., & Basilone, G. (2015). Energy acquisition and allocation to egg production in relation to fish reproductive strategies. Fish Fish., 16(1), 23–57.

56. Middlemiss, K. L., Cook, D. G., Jerrett, A. R., & Davison, W. (2018). Effects of group size on school structure and behaviour in yellow-eyed mullet *Aldrichetta forsteri*. J. Fish Biol, 92(5), 1255–1272.

57. Mukherjee, I., & Bhat, A (2025). Intra- and inter-species information transfer aids tropical fish shoals in detecting predators and food sources. Behav. Ecol. Sociobiol.,79, (105). 10.1007/s00265-025-03651-y

58. Mukherjee, I., & Bhat, A. (2024). The impact of predators and vegetation on shoaling in wild zebrafish. R. Soc. Open Sci., 11(9), 240760.

59. Norin, T., & Clark, T. D. (2016). Measurement and relevance of maximum metabolic rate in fishes. J. Fish Biol., 88(1), 122–151.

60. Norin, T., & Metcalfe, N. B. (2019). Ecological and evolutionary consequences of metabolic rate plasticity in response to environmental change. Phil. Trans. R. Soc. B, 374(1768), 20180180.

61. Peterson, C. H. (1976). Cruising speed during migration of the striped mullet (*Mugil cephalus* L.): An evolutionary response to predation? Evolution, 30(2), 393–396.

62. Pinheiro, J. V., Albuquerque, A., Ferreira, D., Gonçalves, D. M., & Giglio, V. J. (2024). Number of individuals, but not habitat complexity, influences the antipredator behavior of an Amazonian floodplain fish. Neotrop. Ichthyol., 22(03), e240044.

63. Radford, A. N., & Ridley, A. R. (2006). Recruitment calling: a novel form of extended parental care in an altricial species. Curr. Biol., 16(17), 1700–1704.

64. Richards, C. E., & Castagna, M. (1976). Distribution, growth, and predation of juvenile white mullet (*Mugil curema*) in oceanside waters of Virginia’s eastern shore. Chesapeake Sci., 17(4), 308–309.

65. Ridgway, S., Dibble, D. S., & Baird, M. (2022). Sights and sounds dolphins, *Tursiops truncatus* preying on native fish of San Diego Bay and offshore in the Pacific Ocean. Plos one, 17(8), e0265382.

66. Rodriguez-Pinto, I. I., Rieucau, G., Handegard, N. O., Boswell, K. M., & Theobald, J. C. (2024). Environmental impact on visual perception modulates behavioral responses of schooling fish to looming predators. J. Exp. Biol., 227(6), jeb246665.

67. Rosenfeld, J., Van Leeuwen, T., Richards, J., & Allen, D. (2015). Relationship between growth and standard metabolic rate: measurement artefacts and implications for habitat use and life-history adaptation in salmonids. J. Anim. Ecol.84(1), 4–20.

68. Sabal, M. C., Boyce, M. S., Charpentier, C. L., Furey, N. B., Luhring, T. M., Martin, H. W., Melnychuk, M.C., Srygley, R.B., Wagner, M.R., Wirsing, A.J., Ydenberg, R.C., & Palkovacs, E. P. (2021). Predation landscapes influence migratory prey ecology and evolution. Trends Ecol. Evol., 36(8), 737–749.

69. Scarabello, M., Heigenhauser, G. J. F., & Wood, C. M. (1992). Gas exchange, metabolite status and excess post-exercise oxygen consumption after repetitive bouts of exhaustive exercise in juvenile rainbow trout. J. Exp.Biol., 167(1), 155–169.

70. Schindelin, J., Arganda-Carreras, I., Frise, E., Kaynig, V., Longair, M., Pietzsch, T., Preibisch, S., Rueden, C., Saalfeld, S., Schmid, B., Tinevez, J., White, D.J., Hartenstein, V., Eliceiri, K., Tomancak. P., & Cardona, A. (2012). Fiji: an open-source platform for biological-image analysis. Nat. Meth., 9(7), 676–682.

71. Shapiro Goldberg, D., Rilov, G., Villéger, S., & Belmaker, J. (2021). Predation cues lead to reduced foraging of invasive *Siganus rivulatus* in the Mediterranean. Front. Mar. Sci., 8, 678848.

72. Sih, A., Bolnick, D. I., Luttbeg, B., Orrock, J. L., Peacor, S. D., Pintor, L. M., E. P., Preisser, J. S., Rehage.. & Vonesh, J. R. (2010). Predator–prey naïveté, antipredator behavior, and the ecology of predator invasions. Oikos, 119(4), 610–621.

73. Sorato, E., Gullett, P. R., Griffith, S. C., & Russell, A. F. (2012). Effects of predation risk on foraging behaviour and group size: adaptations in a social cooperative species. Anim. Behav., 84(4), 823–834.

74. Soriano-Redondo, A., Franco, A. M., Acácio, M., Payo-Payo, A., Martins, B. H., Moreira, F., & Catry, I. (2023). Fitness, behavioral, and energetic trade-offs of different migratory strategies in a partially migratory species. Ecology, 104(10), e4151.

75. Treherne, J. E., & Foster, W. A. (1981). Group transmission of predator avoidance behaviour in a marine insect: the Trafalgar effect. Anim. Behav., 29(3), 911–917.

76. Whitfield, A. K., Panfili, J., & Durand, J. D. (2012). A global review of the cosmopolitan flathead mullet *Mugil cephalus* Linnaeus 1758 (Teleostei: Mugilidae), with emphasis on the biology, genetics, ecology and fisheries aspects of this apparent species complex. Rev. Fish Biol. Fish., 22(3), 641–681.

77. Wilgers, D. J., Wickwire, D., & Hebets, E. A. (2014). Detection of predator cues alters mating tactics in male wolf spiders. Behav., 151(5), 573–590.

78. Xu, D. D., Wang, C. H., Bi, J. Q., Luo, H., Fu, S. J., Li, B., & Zeng, L. Q. (2024). Physiological and behavioral responses to social isolation and starvation in a social fish. Appl. Anim. Behav. Sci.278, 106384.

79. Zhang, Y., & Lauder, G. V. (2024). Energy conservation by collective movement in schooling fish. eLife 12. *RP90352. doi*, *10*.

80. Zhang, Y., Claireaux, G., Takle, H., Jørgensen, S. M., & Farrell, A. P. (2018). A three-phase excess post-exercise oxygen consumption in Atlantic salmon *Salmo salar* and its response to exercise training. J. Fish Biol. 92(5), 1385–1403.

81. Zhang, W., Cao, Z. D., Peng, J. L., Chen, B. J., & Fu, S. J. (2010). The effects of dissolved oxygen level on the metabolic interaction between digestion and locomotion in juvenile southern catfish (Silurus meridionalis Chen). Comp. Biochem. Physiol. A Mol. Integr. Physiol., 157(3), 212–219.

82. Zimmermann, U., & Curio, E. (1988). Two conflicting needs affecting predator mobbing by great tits, Parus major. Anim.Behav., 36(3), 926–932.

