## Supplementary FIle for "Energetics and behavior during predation in wild, schooling white mullet (*Mugil curema*)"

**Figure S1**

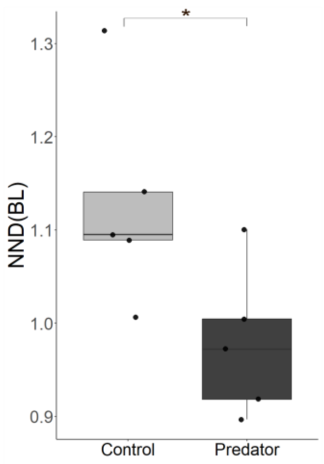


**Figure S1. Lower nearest neighbor distance in predation in groups.** Box and whisker plots showing predator-exposed fish have significantly reduced NND (dark gray) than Control fish (light gray, $p < 0.05$). Each dot denotes escape the NND value for a single school. * indicates a statistically significant difference..

**S2.**

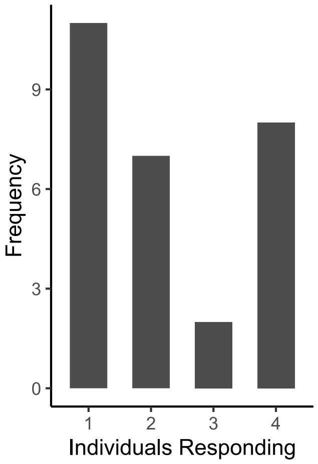


**Figure S2. Bar plot showing the frequency of escape responses by one to four individuals following predator stimuli.** Simultaneous escape by all four individuals occurs relatively infrequently compared to partial group responses (one to three individuals escaping), indicating that escape behavior is not fully synchronized.
